# A photosynthetic checkpoint halts *Synechocystis* sp. PCC 6803 response to nitrogen deprivation during glucose-mediated metabolic imbalance

**DOI:** 10.1101/2025.07.30.667207

**Authors:** Pablo Ortega-Martínez, Joaquín Giner-Lamia, Laura T. Wey, M. Isabel Muro-Pastor, Francisco J. Florencio, Sandra Díaz-Troya

**Affiliations:** Instituto de Bioquímica Vegetal y Fotosíntesis, Universidad de Sevilla-Consejo Superior de Investigaciones Científicas, Américo Vespucio 49, Sevilla, 41092, Spain; Departamento de Bioquímica Vegetal y Biología Molecular, Facultad de Biología, Universidad de Sevilla, Profesor García González s/n, Sevilla, 41012, Spain; Department of Life Technologies, University of Turku, Turku, Finland

**Keywords:** Cyanobacteria, Chlorosis, Nitrogen metabolism, Carbon metabolism, Photosynthesis, Photomixotrophy

## Abstract

Cyanobacteria adapt to nitrogen starvation by undergoing chlorosis, a regulated bleaching process involving the degradation of phycobilisomes, the light harvesting antennae complexes, and accumulation of glycogen. While this response is well-characterized under photoautotrophic conditions, its modulation by external organic carbon sources such as glucose remains poorly characterized. Here, we investigated how glucose affects chlorosis onset in the model cyanobacterium *Synechocystis* sp. PCC 6803. Using an integrative approach combining physiological assays, targeted metabolomics, RNA-sequencing, chlorophyll fluorescence and absorbance spectroscopy, we studied the underlying regulatory mechanisms, focusing on redox control and photosynthetic electron transport. Glucose supplementation prevented bleaching, even when added after nitrogen deprivation symptoms had begun.

This effect was associated with excess glycogen accumulation, disrupted carbon partitioning, and buildup of metabolic intermediates, indicating a metabolic overflow. Despite these physiological differences, transcriptomic responses to nitrogen deprivation were largely similar regardless of glucose supplementation, suggesting regulation at the post-transcriptional or metabolic level. Glucose also impaired photosynthetic electron transport by creating a redox bottleneck at the photosystem II (PSII) acceptor side, leading to decreased electron transport to photosystem I (PSI) and oxidation of the P700 pool. These findings suggest that reduction of the P700 acceptor side is required to trigger chlorosis. Our results demonstrate that glucose uncouples nitrogen sensing from the bleaching process by altering redox balance and photosynthetic electron flow. We propose the existence of a redox-sensitive checkpoint that integrates metabolic state with photosynthetic performance, offering new insights into stress adaptation in cyanobacteria.

## Introduction

Cyanobacteria are an ancient and diverse lineage of oxygenic photosynthetic prokaryotes that have shaped Earth’s biosphere for over 2.5 billion years (Schirrmeister et al. 2016). Throughout their evolutionary history, they have adapted to a wide range of ecological niches by developing diverse physiological and metabolic acclimation strategies to cope with fluctuating environmental conditions. Some cyanobacterial species, including *Synechocystis* sp. PCC 6803 (hereafter *Synechocystis*), possess metabolic flexibility that enables them to grow under photomixotrophic conditions by fixing CO₂ via photosynthesis while simultaneously assimilating exogenous organic carbon sources such as glucose (Kaplan et al. 2008).

As a non-diazotrophic cyanobacterium, *Synechocystis* is unable to fix atmospheric nitrogen (N_2_), and has developed complex regulatory responses to survive the limitation of this essential nutrient. Upon limitation of combined nitrogen forms, such as ammonium (NH₄⁺), nitrate (NO₃⁻), or urea, *Synechocystis* undergoes an adaptive program known as chlorosis (Forchhammer and Schwarz 2019). This process involves a profound reorganization of the photosynthetic machinery and cell metabolism allowing cells to enter a dormant-like state and survive for long periods of nitrogen starvation.

The response to nitrogen depletion is triggered by an imbalance in C/N metabolism (Forchhammer and Selim 2020). In the absence of nitrogen assimilation, the GS-GOGAT cycle becomes limited, which leads to an increase in GOGAT’s substrate 2-oxoglutarate (2-OG), the key C/N sensing metabolite. The accumulation of 2-OG is detected by the signal transduction protein PII, and the global nitrogen regulator NtcA (Muro-Pastor et al. 2001; Esteves-Ferreira et al. 2018; Forchhammer and Schwarz 2019). These proteins coordinate the transcriptomic and metabolic response for the acclimation to nitrogen starvation (Esteves-Ferreira et al. 2018).

A key manifestation in this response is the degradation of the phycobilisomes (PBS), the light harvesting antennae complexes, shifting the cell’s blue-green colouring to orange-yellow, a process termed bleaching. This is a strategy with a dual function. Firstly, it allows the recycling of the nitrogen present in these large protein complexes as a supply for *de novo* synthesis of proteins to adapt to this new nutrient situation. Secondly, the decrease in light-harvesting capacity lowers excitation pressure on the photosynthetic electron transport chain (PETC), thus avoiding potential overreduction and photodamage (Salomon et al. 2013; Baier et al. 2014; Levi et al. 2018; Forchhammer and Schwarz 2019). As an outcome, glycogen reserves are highly increased with the carbon from the phycobiliprotein degradation and the fixed CO_2_ (Forchhammer and Schwarz 2019). This is promoted by the redirection of the carbon flux towards gluconeogenic reactions by the inactivation of phosphoglycerate mutase (PGAM). This regulation is caused by the association of PGAM with the small protein CfrA/PirC, an interactor of PII that becomes highly expressed and released from PII under nitrogen deprivation (Muro-Pastor et al. 2020; Orthwein T et al. 2021).

PBS degradation is an orchestrated process involving several components. Among the most relevant are the <u>n</u>on-<u>bl</u>eaching (Nbl) proteins NblA, NblB and NblD, whose expression is induced by NtcA (Baier et al. 2014; Forchhammer and Schwarz 2019; Krauspe et al. 2021). In *Synechocystis*, the NblA1/NblA2 heterodimer interacts with the PBS (Sendersky et al. 2015; Nguyen et al. 2017), facilitating pigment disassociation by the NblB bilin lyase (Levi et al. 2018) and PBS degradation by the ATP-dependent ClpC-ClpP1-ClpR protease complex (Karradt et al. 2008; Baier et al. 2014). NblD also plays an important role in PBS degradation, although its precise function remains unclear (Krauspe et al. 2021). The process begins from the phycocyanin rods progressing to the allophycocyanin core until nearly complete PBS degradation (Sendersky et al. 2015).

Several scenarios prevent the bleaching process, including limitations on cell growth (e.g. carbenicillin and cerulenin) or photosynthesis imposed by light restrictions or inhibitors (e.g. DCMU and DBMIB) (Salomon et al. 2013; Yoshihara and Kobayashi 2022) or the incorporation of 2-OG through a 2-OG permease (Hickman et al. 2013). Despite being able to grow photomixotrophically, a previous report has observed bleaching inhibition by glucose supplementation in *Synechocystis* (Elmorjani and Herdman 1987).

Additionally, chlorosis upon nitrogen depletion is hindered in specific mutant strains. As expected, mutants defective in the regulation of the response to nitrogen deprivation (e.g. Δ*ntcA* (Sauer et al. 1999)) or in genes required to degrade PBS (Δ*nblA*, Δ*nblB* or Δ*nblD;* (Baier et al. 2014; Levi et al. 2018)) are unable to perform bleaching. An additional group of mutants that fails to perform bleaching are those unable to accumulate glycogen. Mutants lacking ADP-glucose pyrophosphorylase (AGP), the enzyme catalysing the first committed step in the synthesis of glycogen, are unable to degrade their phycobiliproteins in response to nitrogen deprivation, despite sensing nitrogen deficiency and activating the corresponding transcriptional program (Gründel et al. 2012; Hickman et al. 2013; Carrieri et al. 2017). Upon nitrogen deprivation, glycogen-deficient mutants also exhibited growth arrest, metabolic overflow, and a progressive and fast closure of PSII reaction centers, decreasing O_2_ evolution despite retaining PBS complexes (Carrieri et al. 2012; Gründel et al. 2012; Ortega-Martínez et al. 2023).

Despite extensive study of nitrogen starvation responses, the biochemical mechanism by which glucose supplementation halts the chlorosis program in *Synechocystis* under nitrogen deprivation remains unresolved. Notably, this response is counterintuitive: it might be expected that the increased C/N ratio caused by exogenous glucose would promote chlorosis. Instead, here we report that the additional carbon input from glucose supplementation results in a metabolic imbalance that impairs photosynthetic electron transport and prevents PBS degradation. Bleaching is blocked even though nitrogen deficiency is perceived by canonical sensors such as the PII protein and the NtcA-regulon is transcriptionally activated. This suggest that additional regulatory layers - potentially linked to photosynthetic activity and redox balance modulate the bleaching process. Therefore, managing cellular carbon pools becomes crucial for an adequate response to nitrogen deprivation. Furthermore, our results help explain why chlorosis is suppressed under various conditions that disrupt carbon flux or photosynthesis, including growth-limiting environments, glycogen metabolism mutants, and low-light or photosynthesis-inhibiting treatments.

## Materials and methods

### Strains and culture conditions

For routine cell culture, *Synechocystis* sp. PCC 6803 strains were cultivated under photoautotrophic conditions in BG11 medium, with 17.6 mM NaNO_3_ as the nitrogen source, and supplemented with 12 mM NaHCO_3_ (referred as BG11C) (Rippka et al. 1979). Cultures were grown at 30 °C under continuous light (4500k LED lights, 50 µmol photons m^−2^ s^−1^) in conical flasks bubbled with a stream of 1% (v/v) CO_2_ in air. *Synechocystis* cultures growth was monitored by optical density measurement at 750 nm in a UV-Vis GENESYS™180 spectrophotometer (ThermoFisher). Whole-cell absorption spectra were recorded in cultures adjusted to 1 OD_750_ using visible light ranging from 400-800 nm. When required, media were supplemented with the required antibiotics (50 µg ml^−1^ nourseothricin).

For nitrogen deprivation, strains were harvested twice by centrifugation (10 min at 7000 g) in nitrate-free BG11C medium (BG11_0_C) and finally resuspended at 1 OD_750_ in BG11_0_C. Cultures were divided and cultured under control (C) conditions or supplemented with 4 mM glucose (G).

### Generation of mutant strains

The ΔG6PDH strain lacking the *slr1843* gene was obtained by transformation of the WT strain with plasmid pSPARK_ΔG6PDH::Nat(+), which allows for the deletion of the complete *slr1843* ORF. This plasmid contains a noursethricin resistance cassette flanked by sequences 511 pb upstream and 650 pb downstream of the *slr1843* ORF. Complete segregation was confirmed by PCR with primers *slr1843* UP 5’ (5’-GGTGGGACTATTACCGGG-3’) and *slr1843* R (5’-GAGATAGCTTTCTGGG-3’).

### Glycogen quantification and glucose consumption determination

Glycogen content was determined as in (Ortega-Martínez et al. 2023). Briefly, glucose released from purified glycogen and glucose content in the medium were measured with a glucose oxidase assay, employing standard curves of known concentrations of amyloglucosidase-digested glycogen and glucose, respectively.

### Glutamine synthetase activity

Measurements of *in situ* glutamine synthetase activity were performed as described in (Merida et al. 1991) using the Mn^2+^-dependent γ-glutamyl-transferase assay in cells permeabilized with alkyltrimethylammonium bromide (MTA). Concentration of the resulting γ-glutamylhydroxamic was determined spectrophotometrically at 500 nm using a molar extinction coefficient of 0.89 mM^-1^·cm^-1^. One U of GS activity corresponds to the amount of enzyme required to produce 1 μmol product per minute.

### Metabolite extraction and targeted metabolomics

Intracellular metabolites were extracted and determined by LC-MS as described in (Ortega-Martínez et al. 2024). Chromatographic separation was performed with a XSELECT HSS XP 150 mm × 2.1 mm × 2.5 μm (Waters) in an Exion HPLC (Sciex) connected to a QTrap 6500+ (Sciex) operating in negative mode.

### RNA extraction and RNA-seq analysis

Cultures (30 OD_750_) were harvested by centrifugation (10 min, 4°C, 5000 rpm) and pellets frozen in liquid nitrogen and stored at -70°C. Cell pellets were resuspended in 450 µl RLT buffer (RNeasy Plant Mini Kit) supplemented with 4.5 µl β-mercaptoethanol, mixed with glass beads and lysed by mechanical disruption using a vortex-Genie2 with a TurboMix Attachment (2 cycles of 2 min). The lysates were centrifuged (15 s, 4 °C, 13000 rpm) and processed with RNeasy Plant Mini Kit (QIAGEN, #74904). To remove genomic DNA contamination, RNA samples were incubated with 4 U of Turbo DNAse (AM1907, Invitrogen) for 1 hour at 37°C and then purified and concentrated using the RNeasy MinElute Cleanup Kit (QIAGEN, #74204). The integrity of the isolated RNA was assessed by agarose gel electrophoresis with the Bioanalyzer® 2100 system (Agilent Technologies).

The RNA-seq libraries were prepared using the Illumina Total RNA Prep with Ribo-Zero Plus kit (Illumina), following the manufacturer’s protocol. Library quality control included assessment of fragment size distribution and integrity with the Bioanalyzer® 2100, and quantification of DNA concentration using the Qubit™ DNA HS Assay Kit (Thermo Fisher Scientific). The resulting DNA libraries were sequenced on an Illumina NovaSeq 6000 SP-200 platform using 2 × 75 bp paired-end reads. The sequencing was performed by the Genomic Service of CABIMER (Seville, Spain). A total of 121 million reads were obtained, with >94.8% of bases achieving a quality score of Q30 or higher.

The sequencing reads were aligned to the *Synechocystis* sp. PCC 6803 reference genome (NCBI Assembly: GCF_000009725.1; RefSeq: NC_000911.1) using BWA-mem2 alignment algorithm from BWA v2.2.1 (Li 2013). Gene-level raw read counts were generated using the HTSeq-count function from the HTSeq framework v0.11.0 (Anders et al. 2015). The differential gene expression analysis was performed using the DESeq2 package from Bioconductor (Love et al. 2014) in the R environment. Genes with an adjusted p-value < 0.05 were considered significantly differentially expressed. Genes were functionally annotated using the CyanoGenes webserver (www.cyanogenes.ciccartuja.es).

### Oxygen evolution

Oxygen evolution was measured with a Clark-type oxygen electrode (Hansatech) at 30°C. Two ml at 5 µg chlorophyll *a* (chla) ml^-1^ dark-adapted cultures were illuminated for 10 min (50 µmol photon m^-2^ s^-1^) with 5 min dark periods before and after light exposure. To prevent carbon limitation, cultures were supplemented with 10 mM of NaHCO_3_ immediately before measurements. When needed, freshly prepared 2,6-Dichloro-1,4-benzoquinone (DCBQ, 0.25 mM) and potassium ferricyanide (2.5 mM) were added to the chamber.

### Chlorophyll fluorescence analysis and Y(II) calculations

Chlorophyll *a* fluorescence was measured by pulse-amplitude-modulation fluorometry with a Dual-PAM-100 (Walz) using 1.5 ml cultures at 5 µg chla ml^-1^ as described in (Ortega-Martínez et al. 2023). The effective quantum yield of PSII [Y(II)] was determined with the formula (Fm’-Fs)/(Fm’) where Fm’ is maximal fluorescence and Fs is basal fluorescence, both measured during exposure to actinic light. Other parameters analyzed were basal fluorescence in the dark (Fo), qL and qP both relate to the redox state of the plastoquinone pool (lake and puddle models, respectively (Kramer et al. 2004)). These parameters were obtained from the recording of an induction curve, exposing dark-adapted cultures to actinic light (40 µmol photon m^-2^ s^-1^) followed by a post-illumination recovery and multiple saturation pulses (5000 µmol photon m^-2^ s^-1^).

### NAD(P)H fluorescence measurements

The NADPH/9-AA module of a DUAL-PAM (Walz, Germany) was used to measure light-induced NADPH redox kinetics on 7.5 µg chla ml^-1^ samples at 30°C by monitoring the changes in fluorescence with excitation at 365 nm and detection between 420 and 580 nm as in (Ortega-Martínez et al. 2024).

### Determination of P700 redox kinetics

P700 redox kinetics were measured by pulse-amplitude-modulation fluorometry with a Dual-PAM-100 (Walz) using intact cells at a concentration of 12.5 μg chla ml^-1^ at room temperature similarly to (Shimakawa and Miyake 2018a) and based on the method of (Klughammer and Schreiber 2008). Before recording, cells were adapted in the dark for 10 min. During an induction curve with actinic light at 80 µmol photons m^-2^ s^-1^, 300 ms saturation pulses (SP) at 5000 µmol photons m^-2^ s^-1^ with peak emission at 635 nm were supplied at the indicated times to calculate P700 parameters: the PSI quantum yield of photochemical energy conversion [Y(I) = (Pm-P)/Pm] and the non-photochemical energy dissipation due to donor-side [Y(ND)= P/Pm] and acceptor-side [Y(NA)= (Pm - Pm)/Pm] limitations (Fig. S1).

### Graphic representation and statistical analysis

In general, visualization of continuous data with the associated standard error of the mean was performed using dplyr and ggplot2 (Tidyverse packages) in R. Figures containing discrete data and statistical analysis were obtained using the software GraphPad Prims 8.0.1. The statistical method used and the number of experimental replicates are described in the figure legends

## Results

### Effect of glucose supplementation in *Synechocystis* under nitrogen deprivation

To address the mechanism by which glucose inhibits the chlorosis process we analysed the physiological response of *Synechocystis* to nitrogen deprivation in the presence of glucose. WT cells, previously grown photoautotrophically in nitrogen-replete medium, were transferred to nitrogen-depleted medium in the absence (C, control) or presence of 4 mM glucose (G, glucose). As expected, while the control culture started chlorosis in response to nitrogen deprivation, glucose supplementation completely blocked bleaching, and the cells maintained their pigmentation until 48 h (Fig 1a), in agreement with previous results (Elmorjani and Herdman 1987).

**Figure 1:**
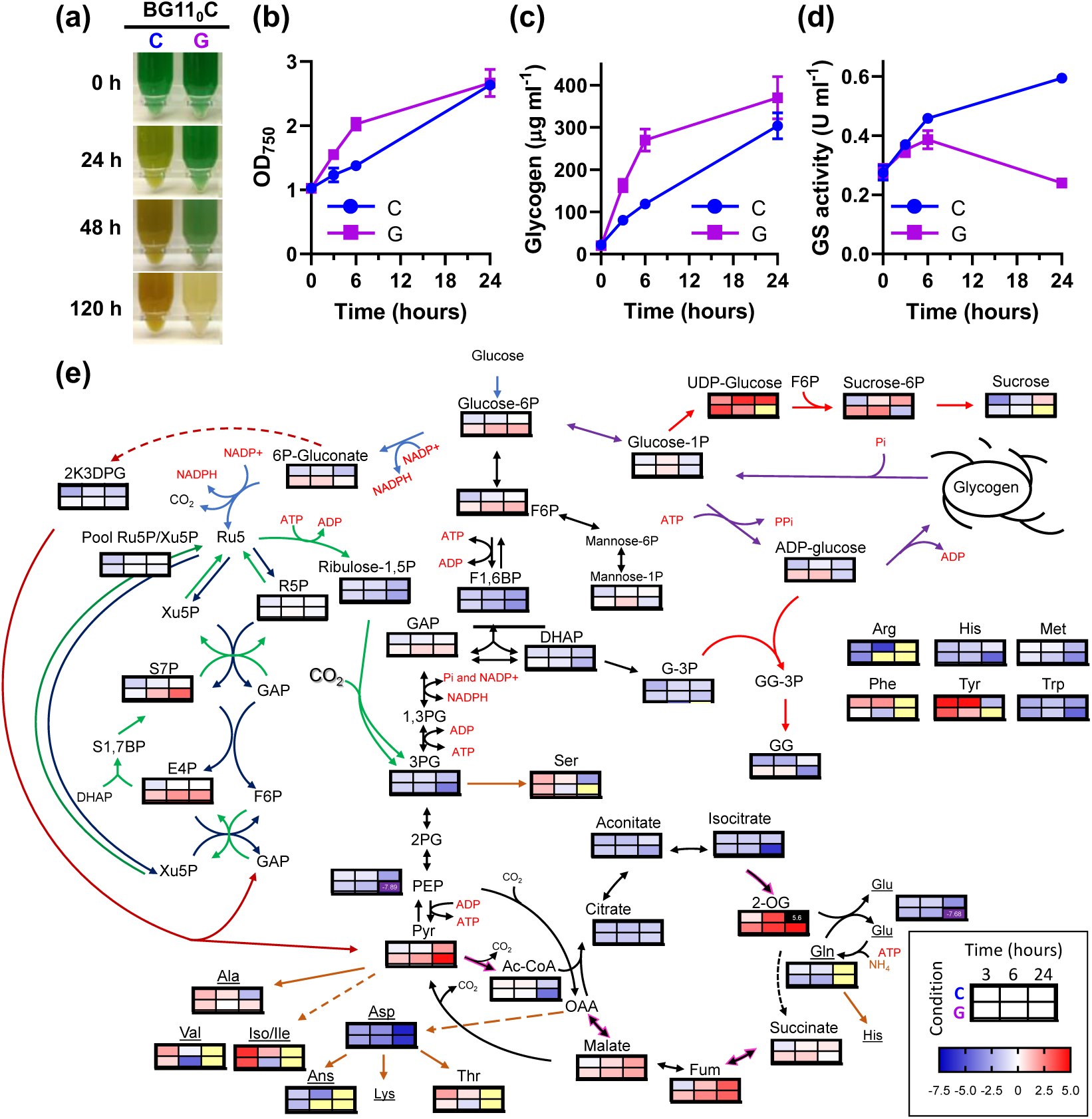
Physiological characterization of the effect of glucose supplementation on the response to nitrogen deprivation in WT *Synechocystis*. WT cells grown photoautotrophically in nitrogen-replete medium were harvested and resuspended in nitrogen-free medium (BG11_0_C) at 1 OD_750_. Culture was divided and grown under control (C) conditions or supplemented with 4 mM of glucose (G). (a) Photographs of the cultures at 0, 24, 48 and 120 hours after the onset of nitrogen limitation. (b) Growth curves measured as OD_750_. (c) Glycogen content. (d) Glutamine synthetase activity expressed as U ml^-1^..(e) Relative abundance of metabolites measured by LC/MS. Individual heatmaps for each metabolite are embedded within a metabolic network diagram representing the log2 fold-change of the mean abundance relative to 0 hours and normalized to OD_750_. Metabolites coloured: not detected (yellow), log2 fold-change > 5 (black), log2 fold-change < 7.5 (purple). Measurements in b-e were performed at a at 0, 3, 6 and 24 hours after onset of nitrogen limitation. Data in (b-d) and (e) represent the mean±SEM of 6 and 4 independent biological replicates, respectively. Abbreviations: F6P (Fructose-6P), F1,6BP (Fructose-1,6-BP), 2K3DPG (2-Keto-3-Deoxy-Phosphogluconate, Ru-5P/Xu-5P (ribulose-5P/xylulose-5P pool), R5P (Ribose-5P), S7P (Sedoheptulose-7P), E4P (Erythrose-4P), GAP (glyceraldehyde-3P), DHAP (dihydroxyacetone-P), 3PG (3P-glycerate), PEP (phosphoenolpyruvate), Pyr (Pyruvate), AcCoA (Acetyl-CoA), 2-OG (2-oxoglutarate), Fum (Fumarate), OAA (oxalacetate), G-3p (glycerol-3P), GG (glucosylglycerol-3P).

Glucose supplementation induced rapid physiological alterations in the response to nitrogen deprivation that ultimately compromised viability. Within the first six hours of nitrogen deprivation, we observed a rapid rise in OD_750_ (OD_750_ from 1 to 2.1 ± 0.25) and glycogen reserves (269 µg ml^-1^); representing a 2.28 fold-change increase in glycogen content compared to the control condition (118 µg ml^-1^) (Fig. 1b-c). This was accompanied by a progressive consumption of glucose from the medium, with approximately 1.2 ± 0.18 mM left in the medium after 24 hours (Fig. S2a).

However, despite this physiological response and maintenance of the phycobiliprotein peak (625 nm), the whole-cell absorption spectra presented a flattening pattern (Fig. S2b) indicative of cell stress and compromised cellular viability. This was further corroborated by the significant increase in reactive oxygen species (ROS) after 24 hours (Fig. S2c), and by the whitening of the cultures after 120 h of incubation (Fig. 1a).

Despite the lack of bleaching, glucose supplemented cells sensed the nitrogen depletion and responded accordingly, as indicated by two key markers of nitrogen deficiency: increased glutamine synthetase (GS) activity and PII hyperphosphorylation (Fig. 1d, S2). These two responses occurred similarly during the first hours of nitrogen deprivation both in the presence and absence of glucose. Ultimately, after 24 hours in the presence of glucose, GS activity was decreased, most likely due to the lethality of this condition, with general protein loss and ROS increase (Fig. S2b-c, S3) (Robles-Rengel et al. 2019).

Metabolomic analysis confirmed the expected increase in the nitrogen status sensor 2-OG, key in the responses of GS activity and PII phosphorylation state, under nitrogen deprived conditions both in the presence and absence of glucose (Fig. 1e and Table S1). However, important alterations were observed in the metabolome of glucose supplemented cells. In contrast to control nitrogen deprived cells, glucose supplementation led to a progressive accumulation of glycolytic and pentose phosphate pathway intermediates, including glucose-6-phosphate (G6P), fructose-6-phosphate (F6P), sedoheptulose-7-phosphate (S7P), erythrose-4-phosphate (E4P), and pyruvate, as well as the glycogen precursor ADP-glucose (Fig. 1e and Table S1).

In the absence of glucose, nitrogen starvation resulted in a transitory increase in some amino acids and a severe depletion of all the analysed pools of amino acids after six to 24 hours (Fig. 1e and Table S1). This depletion occurred earlier in the presence of glucose, with minimal levels of glutamine, threonine, valine, phenylalanine or arginine detected after the first three hours. Notably, methionine, an essential initiator of protein synthesis, was nearly undetectable after 24 hours (Fig. 1e and Table S1).

Interestingly, the observed increase in G6P and 6PG, along with the subsequent generation of reducing power (NADPH) through the oxidative pentose phosphate (OPP) shunt, was proposed as a potential key factor in preventing chlorosis (Gründel et al. 2012). To test this hypothesis, we disrupted the OPP shunt by generating a mutant strain lacking glucose-6-phosphate dehydrogenase (G6PDH). The WT and the ΔG6PDH mutant strain were subjected to nitrogen depletion under control or glucose supplementation conditions (Fig. 2). Like the WT, the ΔG6PDH strain could not undergo bleaching when supplemented with glucose, losing the chlorophyll pigments within 48 hours (Fig. 2a) despite exhibiting undetectable 6PG levels (Fig. 2b). In fact, while the WT maintained the typical NAD(P)H light-dependent synthesis trace even 24 hours after nitrogen depletion, the presence of glucose caused an immediate change in the NAD(P)H kinetics with no increase in its associated fluorescence (Fig. 2c), resembling the similar effect observed in the control (Ortega-Martínez et al. 2024). However, as expected due to the impediment of NADPH synthesis by the OPP shunt, the ΔG6PDH strain presented light-dependent increase in the NAD(P)H fluorescence kinetics in the presence of glucose, although the trace is slightly affected compared to the control condition (Fig. 2c). This suggests that 6PG accumulation and potential overreduction of the NADPH pool are not the cause behind the lack of bleaching. Interestingly, while disrupting the OPP shunt resulted in mild alterations in metabolites related to the glucose-6-P crossroads, it did not impact the bleaching process (Fig. 2b). These effects of the ΔG6PDH on the NAD(P)H fluorescence kinetics and metabolic alterations were also observed in previous studies (Maruyama et al. 2019; Hatano et al. 2022).

**Figure 2:**
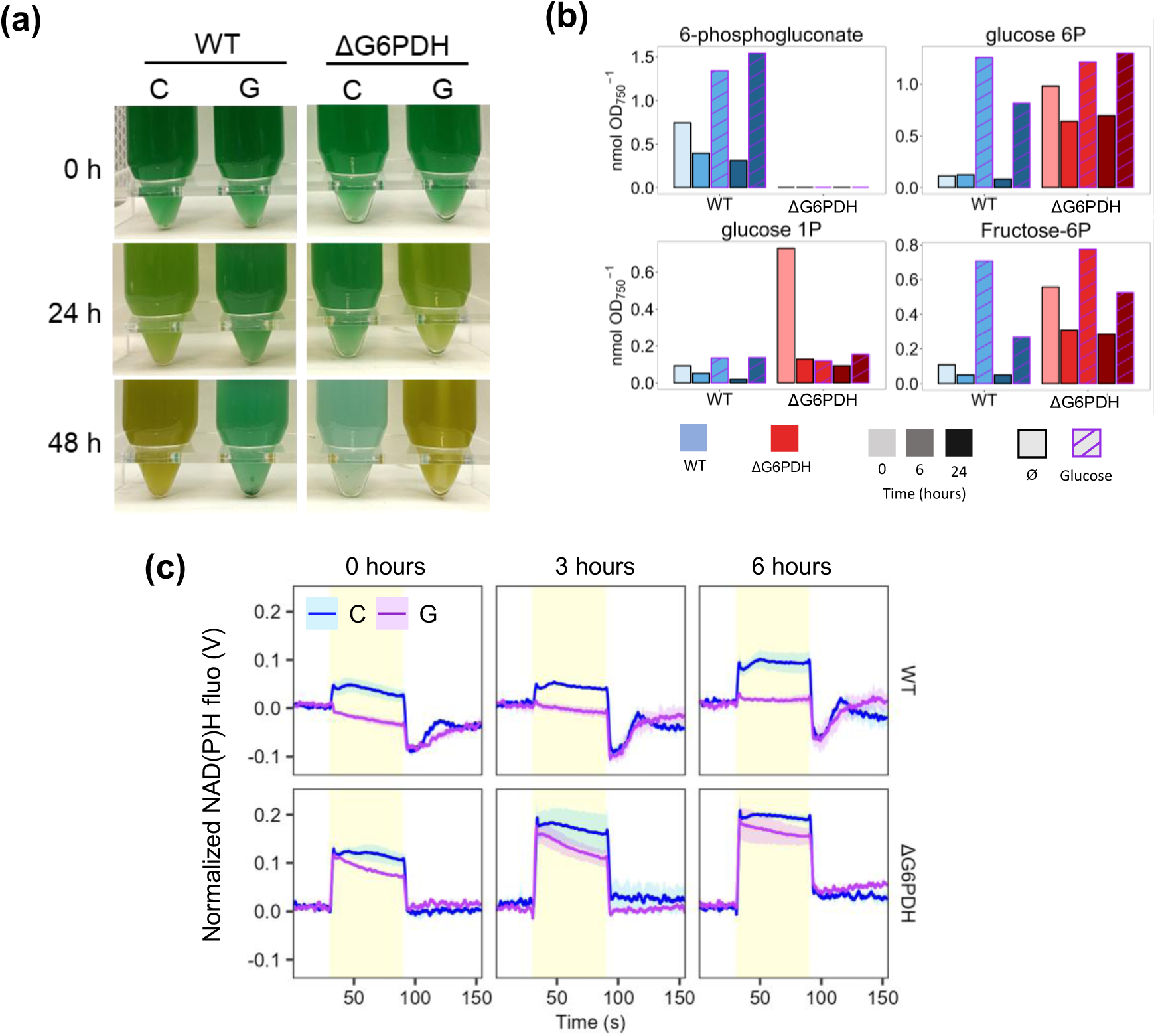
Effect of glucose supplementation on the response to nitrogen deprivation of the ΔG6PDH mutant, lacking gluocose-6P dehydrogenase. WT and ΔG6PDH strains grown in nitrogen-replete medium were harvested and resuspended in nitrogen-free (BG11_0_C) medium at 1 OD_750_. Cultures were divided and grown under control (C) conditions or supplemented with 4 mM of glucose (G). (a) Photographs of the cultures at 0, 24 and 48 hours after the onset of the experiment. (b) Measurements of metabolites from the first steps in glucose assimilation by LC/MS at 0, 6 and 24 hours. (c) NAD(P)H fluorescence light-dependent kinetics of WT and ΔG6PDH cultures (12.5 μg chl mL^-1^) at 0, 3 and 6 hours after the onset of the experiment Data in (b) are from one experiment; data in (c) represent the mean±SEM of 3 independent biological replicates

Overall, these results indicate that glucose intake in nitrogen deprived medium fuels glycogen reserves. However, achieving high glycogen levels early during nitrogen depletion might exceed its total storage capacity, leading to a metabolic overflow like what is observed when glycogen synthesis is limited and thus causing cell death.

### Comparative transcriptome changes induced by glucose upon nitrogen depletion

As the phosphorylation state of PII indicated that glucose did not alter nitrogen sensing, we next proceed to analyse its effect on the transcriptomic response by RNA-seq. *Synechocystis* grown photoautotrophically in nitrogen-replete medium (0 h) were transferred to nitrogen-depleted media, either without (C, control) or with 4 mM glucose (G, glucose), and sampled after three and six hours. A total of 3228 transcripts were detected (Table S2). At the 3-hour time point, we observed a comparable number of differentially expressed genes (DEGs; fold change > 1.5, adjusted *p* < 0.05) in both conditions, with 428 DEGs in Control and 575 DEGs in Glucose compared with nitrogen replete sample (0 h) (Table S3). By six hours under nitrogen deprivation, the transcriptional responses began to diverge more markedly, with 480 DEGs in the control condition and 921 DEGs in the glucose-supplemented condition.

Despite these differences, the global transcriptomic responses to nitrogen starvation in the presence and absence of glucose remained highly correlated across all genes, particularly at three hours (Pearson correlation coefficient: 0.92) and to a lesser extent at six hours (Pearson correlation coefficient: 0.79) (Fig. 3a). When focusing specifically on genes associated with the chlorosis program (230 genes related to photosynthesis and respiration, carbohydrate and carbon central metabolism and nitrogen assimilation and metabolism (Table S4)), we observed that these genes consistently exhibited similar patterns of differential expression in both treatments (Fig. 3b, S3). This indicates that the observed transcriptomic divergence is largely driven by other genes not directly linked, according to current literature, to chlorosis-related cellular processes. Notably, this divergence was more pronounced at six hours, where a substantial proportion of genes showing differential expression patterns between conditions were annotated as hypothetical or of unknown function (NC in Fig. 3a), suggesting that glucose availability may importantly affect the expression of poorly characterized regulatory or metabolic pathways.

**Figure 3:**
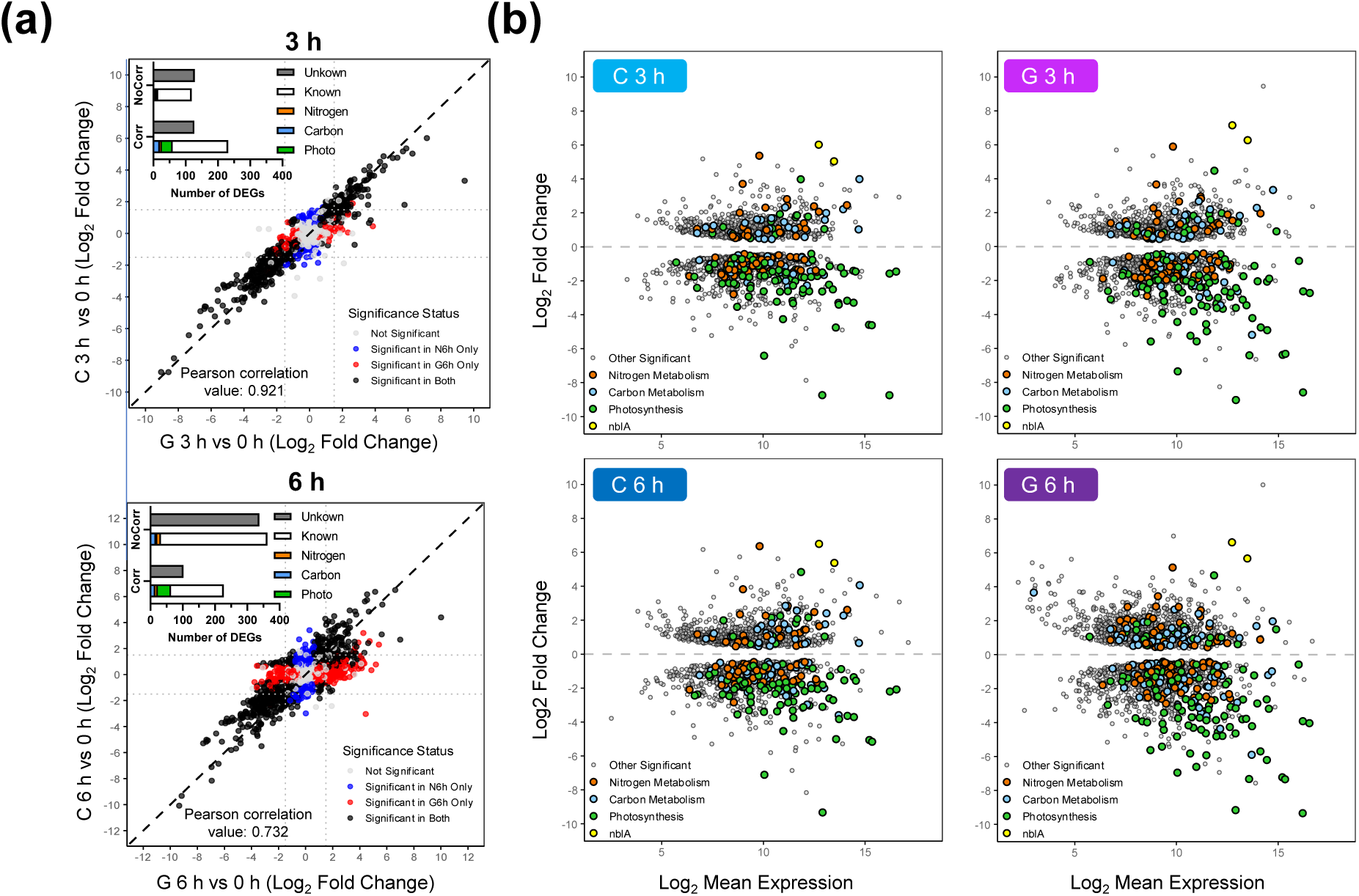
Effect of glucose supplementation on the transcriptomic response of *Synechocystis* to nitrogen deprivation. WT cells grown photoautotrophically in nitrogen-replete medium (0 h) were harvested and resuspended in nitrogen-free medium (BG11_0_C) at 1 OD_750_. Culture was divided and grown under control (C) conditions or supplemented with 4 mM of glucose (G). (a) Scatter plots showing the correlation of gene expression between C and G conditions after 3 and 6 hours of nitrogen deprivation. Pearson correlation coefficients are indicated at the bottom of each plot. Functional annotation of known and unknown differentially expressed genes (DEGs) that showed correlation (Corr) or no correlation (NoCorr) is summarized in the top-left insets. Among the annotated genes, those involved in carbon metabolism (Carbon), photosynthesis and respiration (Photo), and nitrogen metabolism (N) are highlighted in blue, green, and orange, respectively. (b) MA plots showing log₂ fold-change versus mean expression for each gene under nitrogen deprivation in control (C) conditions or supplemented with glucose (G) after 3 and 6 hours of treatment, compared to initial nitrogen-replete state (0 h). The contrasts shown are: C 3 h vs 0 h, C 6 h vs 0 h, G 3 h vs 0 h, and G 6 h vs 0 h. All genes represented (dots) show significant differential expression (adjusted p-value < 0.05). Genes annotated with functional categories related to nitrogen deprivation response are highlighted as coloured dots: carbon metabolism (blue), photosynthesis and respiration (green), nitrogen metabolism and assimilation (orange), and *nblA* genes (yellow). The remaining significant genes are shown in grey.

The well characterised transcriptional response to nitrogen deprivation (Krasikov et al. 2012; Matthias et al. 2014; Carrieri et al. 2017; Giner-Lamia et al. 2017; Esteves-Ferreira et al. 2018) occurred regardless of glucose presence. This common response includes (Fig 3): (i) Upregulation of nitrogen uptake systems, including those for ammonium (*amt*), urea (*urt*), and nitrate (*nrtD, sll1082*). (ii) Adjustment of nitrogen metabolism, with upregulation of *glnN* (*slr0288*), encoding GSIII, and downregulation of genes encoding GS inactivating factors IF7 (*gifA, ssl1911*) and IF17 (*gifB, sll1515*), in agreement with GS activity measurements (Fig. 1d). This also included repression of the genes encoding the ferredoxin-nitrite reductase (*nirA, slr0898*), and the PII protein interactor A (*pirA, ssr0692*), mediating arginine synthesis. (iii) Coordination with carbon metabolism, with strong upregulation of *cfrA/pirC* (*sll0944*), a nitrogen-responsive metabolic regulator controlling carbon flow (Muro-Pastor et al. 2020). Other transcriptional changes related to the regulation of carbon metabolims were dowregulation of genes coding for ADP-glucose pyrophosphorylase (*agp*, *slr1176*), phosphoribulokinase (*prk*, *sll1525*), and both RuBisCO subunits (*rbcS, slr0012* and *rbcL, slr0009*). (iv) Repression of photosynthesis-related genes, including genes coding for phycobiliproteins such as allophycocyanin (APC genes) and phycocyanin (cpcA-D, G) chains, subunits or linkers, photosystems (psa and psb subunits), several ATP synthase subunits (atpA-I), or proteins of the carbon dioxide concentrating mechanism (CCM) (*sll1029*, *sll1030*). In contrast, genes encoding subunits of NADH dehydrogenase (*slr0851*), and cytochrome oxidases (*cyd*, *slr1380*; *slr1379*; *cta, slr1136*; *sll1899*) were upregulated. (v) Finally, of special interest is the strong upregulation of *nblA* genes (*ssl0452* and *ssl0453*), required for phycobiliproteins degradation. These were among the most upregulated genes (log_2_ FC > 5) in both conditions and times tested, indicating that the transcriptional response of phycobiliprotein degradation was promoted (Table S4 and Fig. S4). It is also important to note that many of these genes belong to the NtcA regulon (Giner-Lamia et al. 2017). Therefore, these data confirm that glucose treatment does not interfere in the transcriptomic response to nitrogen deprivation, and specifically with the NtcA-mediated response.

However, glucose supplementation also led to differential transcript changes (Table S4 and Fig. S4). After three hours, only 46 genes presented statistically significant differences with most of them of unknown function. However, the two genes most highly induced by glucose were *fabG* (*sll0330*), implicated in fatty acid biosynthesis and considered as potential PhaB (acetoacetyl-CoA reductase) (Zhang et al. 2017) and the PGR5-like homolog *ssr2016*. These two genes were also upregulated in the absence of glucose, but to a lesser extent. Another DEG in glucose is the Site-2-Protease *sll0528*, related to carbon/nitrogen homeostasis and stress response (Lei et al. 2014; Lin et al. 2025), being highly expressed compared with the control condition (log_2_ FC of 2.3 and 3.91 after three and six hours). In addition, several heat shock (*hsp17, sll1514* and *htpG*, *sll0430*) and high light inducible genes (*ssr2595* and *ssl2542*) were among the most notorious changes detected. Conversely, several genes related to the CCM and PBS assembly were specially repressed in the presence of glucose. Some central carbon metabolism genes such as those encoding pentose-5-phosphate-3-epimerase (*sll0807*), phosphoglucomutase (*sll0726*), phosphoglycerate mutase (*slr1124*) and acetyl-coenzyme A synthetase (*sll0542)* presented a specific depleted expression with glucose. After six hours, an additional set of genes were also induced, mainly with unknown function or related to stress response.

### Photosynthetic characterization during nitrogen deprivation and glucose supplementation

Prior research has demonstrated that metabolic overflow (specifically because of glycogen synthesis deficiency) impairs photosynthesis (Holland et al. 2016; Ortega-Martínez et al. 2023), which can block the bleaching process (Yoshihara and Kobayashi 2022). To evaluate the impact of glucose on photosynthetic performance during nitrogen deprivation, we analysed oxygen evolution in cultures transferred to nitrogen-depleted medium in the absence (C, control) or presence of 4 mM glucose (G, glucose). (Fig. 4, 5). In the control, the O_2_ evolution of the cultures retained c. a. 80±9% and 40±5% of the initial rate by six and 24 hours, respectively (Fig. 4a). In contrast, glucose supplementation decreased O_2_ evolution, reaching 15±10% of the initial rate by six hours and becoming abolished by 24 h. However, this impairment was alleviated by adding the artificial PSII electron acceptor DCBQ, indicating that the oxygen evolving complex of PSII remained functional after six hours of nitrogen deprivation in glucose supplemented cultures. (Fig. S5).

**Figure 4:**
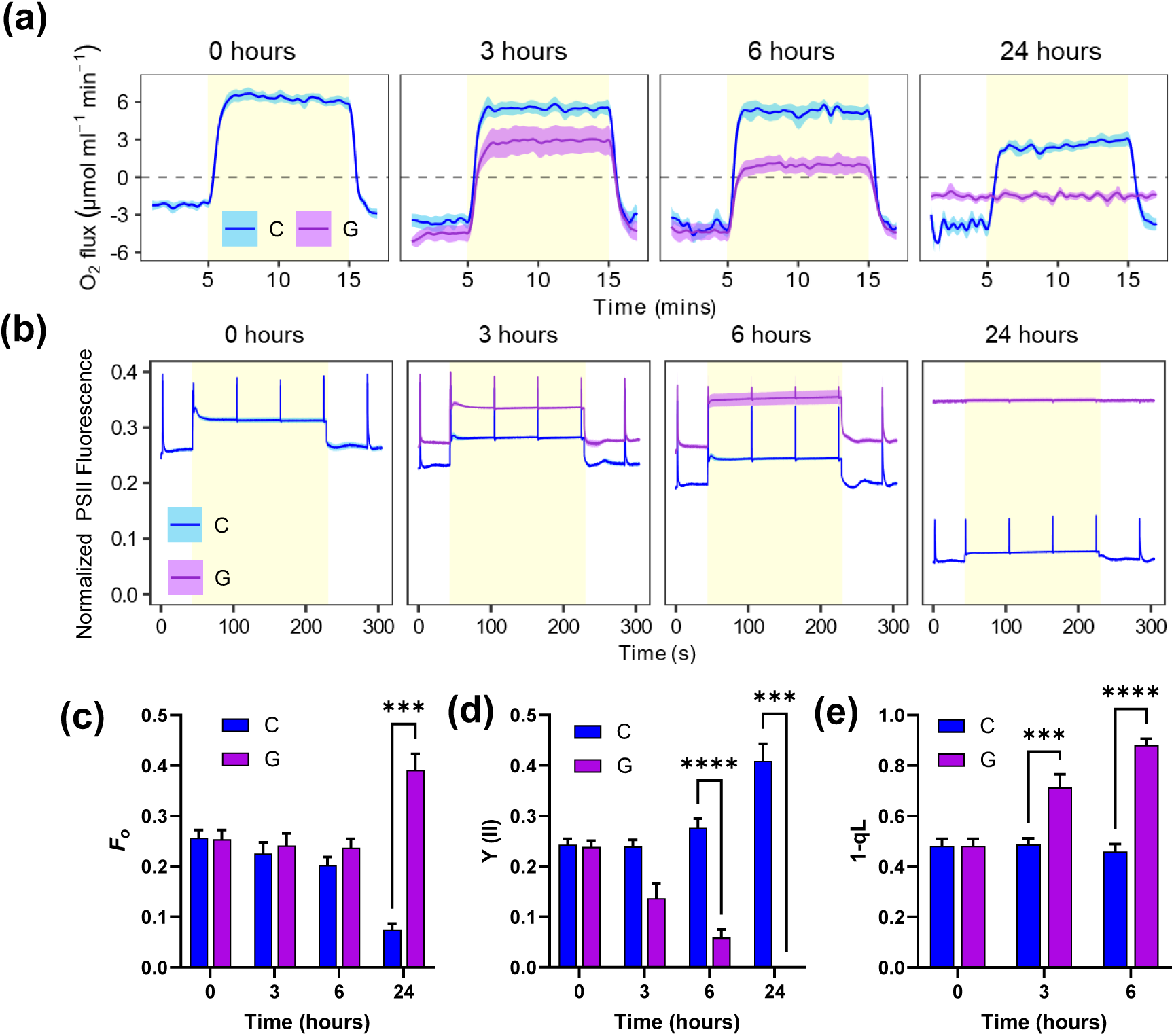
PSII characterization during the response to nitrogen deprivation with glucose supplementation in WT *Synechocystis*. WT cells grown in nitrogen replete medium (BG11C) were harvested and resuspended in nitrogen-free medium (BG11_0_C) at 1 OD_750_. The culture was divided and grown under control (C) conditions or supplemented with 4 mM of glucose (G). (a) Oxygen evolution rates of cultures (5 μg chl mL^-1^) at 0, 3, 6 and 24 hours time points after nitrogen deprivation illuminated with 50 μmol photons m^−2^ s^−1^. (b) Chlorophyll fluorescence induction curves of C and G cultures at timepoints 0, 3, 6 and 24 hours after nitrogen deprivation. Fluorescence values were normalized to the min value of fluorescence trace. Parameters obtained from the traces of chlorophyll fluorescence induction curves are summarized in (c) basal fluorescence in the dark (*F_o_*), (d) effective quantum yield of PSII Y(II), and (e) 1-qL estimations. Data represent the mean±SEM of 5 independent biological replicates.

Chlorophyll fluorescence analysis revealed a progressive decrease in dark-adapted basal fluorescence (*F_o_*) during nitrogen depletion for the control condition under photoautotrophic conditions (Fig. 4b), likely due to phycobiliprotein degradation (Fig. S2b). The effective quantum yield of PSII [Y(II)] under moderate actinic light remained stable during the first 6 h of nitrogen depletion, consistent with the O_2_ evolution (Fig. 4a), but increased to ∼ 0.4 by 24 h (Fig 4b). This rise likely reflects lower excitation energy transfer to PSII due to PBS degradation (Fig. S2b), which lowers both Fs and Fm’, leading to an apparent increase in Y(II) despite some decline in O_2_ evolution (Fig. 4a). In contrast, glucose treatment increased Fs but no Fm’, without detectable decrease in *F_o_* (Fig. 4c). Consequently, there was a severe decrease in Y(II), reaching values as low as 0.05±0.016 after six hours, which is 18% of the value of the control condition at that time point (Fig. 4d). After 24 hours, cells displayed no response to actinic light or saturating pulses, consistent with protein degradation and cell death.

Analysis of the coefficient of photochemical quenching based on the lake model (qL) of antennae connectivity revealed progressive PSII reaction centre closure and increased Q_A_ reduction, suggesting an increasingly reduced plastoquinone (PQ) pool (Fig. 4e). Similar trends were observed using the pool model estimation (1-qP) (Fig. S6). An unchanging 1-qL parameter during 6 h of nitrogen depletion under control condition indicated unchanged redox state of the PQ pool.

To elucidate the impact of this altered photosynthetic capacity on PSI performance, we evaluated P700 redox kinetics with light induction curves in a Dual-PAM-100 (Fig. 5). As a control for PSI acceptor side limitation, we included treatment with 5 mM glycolaldehyde (Gla), an inhibitor of phosphoribulokinase that arrests the CCB cycle by preventing RuBisCO substrate regeneration. Similar to other treatments that interfere with photosynthesis, Gla treatment prevented the bleaching process causing the culture to turn blue after 24 hours of nitrogen depletion (Fig. 5a). The NAD(P)H kinetics showed a constant increase in NAD(P)H fluorescence during the light period (Fig. S7) instead of achieving a steady state, as observed in other conditions (Fig. 2), which confirms the effectiveness of the treatment.

**Figure 5:**
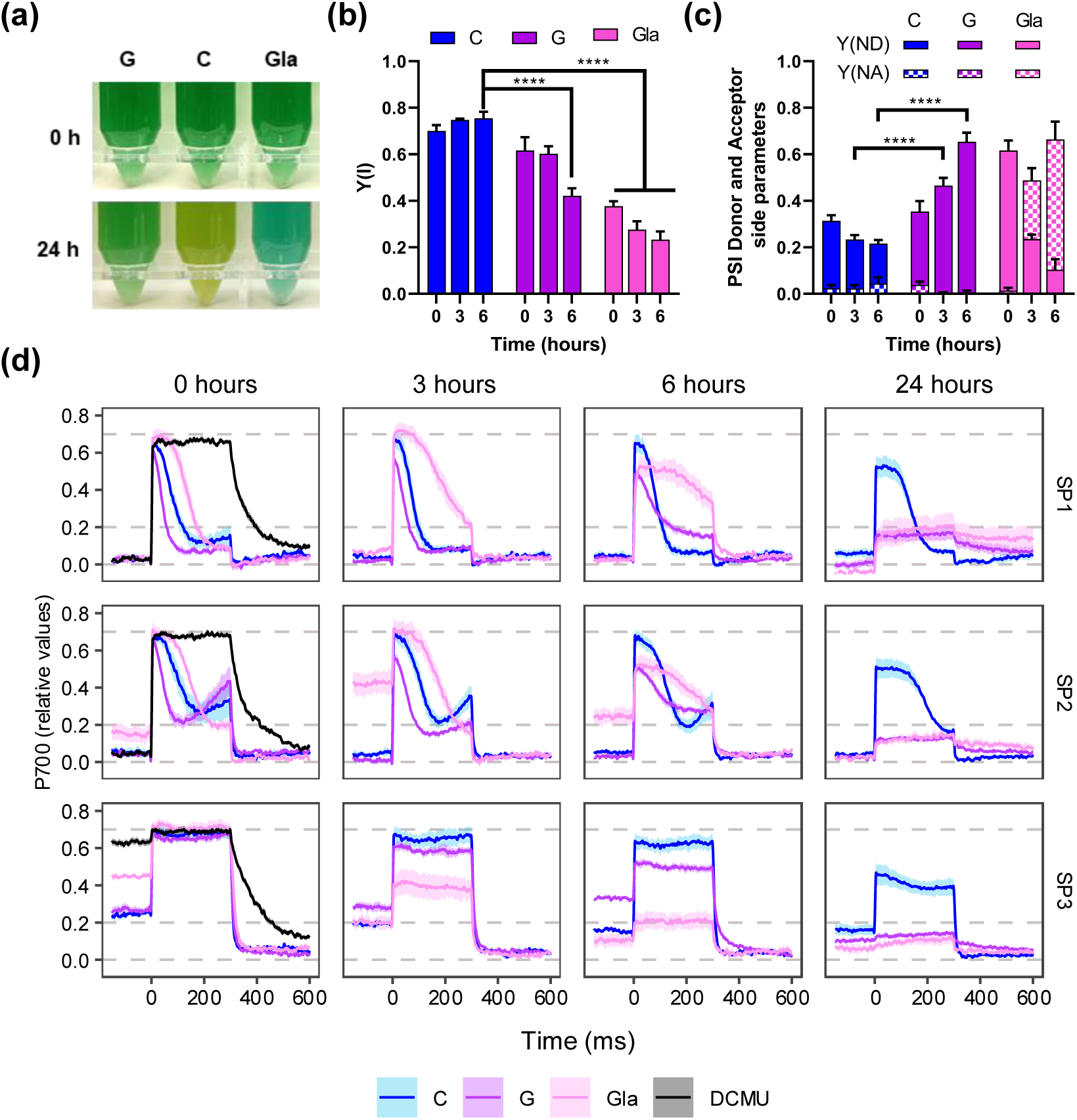
P700 characterization during the response to nitrogen deprivation with glucose and glycolaldehyde supplementation in WT *Synechocystis*. WT cells grown in nitrogen replete medium (BG11C) were harvested and resuspended in nitrogen-free medium (BG11_0_C) at 1 OD_750_. The culture was divided and grown under control (C) conditions or supplemented with 4 mM of glucose (G) or 5 mM of glycolaldehyde (Gla). (a) Photographs of the cultures at 0 and 24 hours. (b) Quantification of PSI effective quantum yield [Y(I)] during the SP3 of the induction curve for all the conditions at 0, 3 and 6 hours after nitrogen depletion. (c) Quantification of PSI donor-side limitation [Y(ND)], and acceptor-side limitation [Y(NA)] during the SP3 of the induction curve for all the conditions at 0, 3 and 6 hours after nitrogen depletion. (d) Analysis of the redox state of P700 by the changes in near-infrared absorbance upon the saturation pulses (SP1-3; see Figure S1) during an induction curve in a Dual-PAM-100 for all the conditions at 0, 3, 6 and 24 hours after nitrogen depletion. For PSII inhibition, 20 µM DCMU was added to control culture at 0 hours. Dashed lines represent from bottom to top: basal oxidation level, the steady-state P700 oxidation level under actinic light (P) and maximum P700 oxidation level (Pm) under a saturation pulse during actinic light of nitrogen replete condition at time 0 h. Data represent the mean±SEM of 4 independent biological replicates. Statistical significance was denoted **** P<0.0001; otherwise indicate no significance (two-way ANOVA).

To evaluate the redox state of PSI, we analyzed the effect of saturating pulses in the dark (SP1), after far-red illumination (SP2) and during actinic illumination (SP3) (Fig. S1). Under control conditions at the onset of nitrogen depletion, P700 presented typical redox kinetics during the SPs (Fig. 5d). In the dark, the SP1 caused a fast oxidation of the fully reduced P700 reaction center to its maximal oxidized state, which then was reduced by electrons from PSII (reduction phase absent in the SP1 and SP2 P700 redox kinetics of DCMU-treated cells). This reduction persisted during the rest of the SP due to limitations on P700 acceptor side primarily due to the lack of actinic light to switch on the CBB cycle (Shimakawa and Miyake 2018b). Under far-red illumination, which preferentially excites PSI, the P700 redox kinetics during SP2 also featured a re-oxidation phase after 150 ms by the action of the flavodiiron proteins (FLVs) (Shimakawa and Miyake 2018a). However, under actinic light, the P700 pool was partially oxidized due to lack of limitation on the acceptor side (Shimakawa and Miyake 2018b). During actinic light, P700 presents a partially oxidized state that becomes fully oxidized during SP3, just to return to a reduced state after the SP. Addition of DCMU caused a delay in the reduction of P700 after the SPs and an almost complete oxidized state during actinic light (SP3) due to the restricted electron donation form PSII (Fig. 5d).

Supplementation of the nitrogen deprived cultures with glucose or Gla had immediate effects in the redox kinetics of P700. Glucose quickened the reduction of P700 during SP1 and SP2, and a stronger re-oxidation associated with the FLVs activities during SP2. Conversely, Gla slowed the reduction of P700 during SP1 and SP2, and had a partially oxidized P700 pool during the actinic light prior to SP3. Notably, Gla treatment also increased the steady-state oxidation level of P700 during far red light suggesting that Gla not only inhibits the CBB cycle but also interferes in cyclic electron transport (CET) as observed previously (Kusama et al. 2022).

Through 3 and 6 h of nitrogen deprivation, control cells maintained their P700 redox kinetics (Fig. 5d). Glucose supplementation progressively increased the oxidized pool of P700 during actinic light (SP3), decreased the maximum oxidisable P700 pool, decreased Y(I) (Fig. 5b), and increased limitation on the P700 donor side [Y(ND)] (Fig. 5c).

At 24 hours of nitrogen deprivation and treatment with glucose or Gla, the P700 photooxidation signal was negligible. In summary, these data suggest a restricted electron flow in the presence of glucose when nitrogen is depleted that results in the oxidation of P700. Therefore, maintaining some of the P700 pool in a reduced state during illumination seems to be critical to perform bleaching.

### Effect of glucose addition at different times during the bleaching process

To determine whether the inhibitory effect of glucose on the bleaching process is restricted to its onset or if it can halt the process after it has been initiated, glucose was added to WT cultures at different times or concentrations after nitrogen removal. Nitrogen depletion response was halted when glucose was added regardless of the stage of the bleaching process, even after phycobiliproteins degradative process had already started (Fig. 6a). This indicates that the inhibitory effect of glucose on the bleaching process is not restricted to the onset of the response to nitrogen deprivation. Interestingly, the response was dose-dependent and reversible: if the glucose provided was below a certain threshold (1 mM glucose) then bleaching proceeded (Fig.6b). This indicates that cells can accommodate certain levels of disturbance in metabolic intermediates levels.

**Figure 6:**
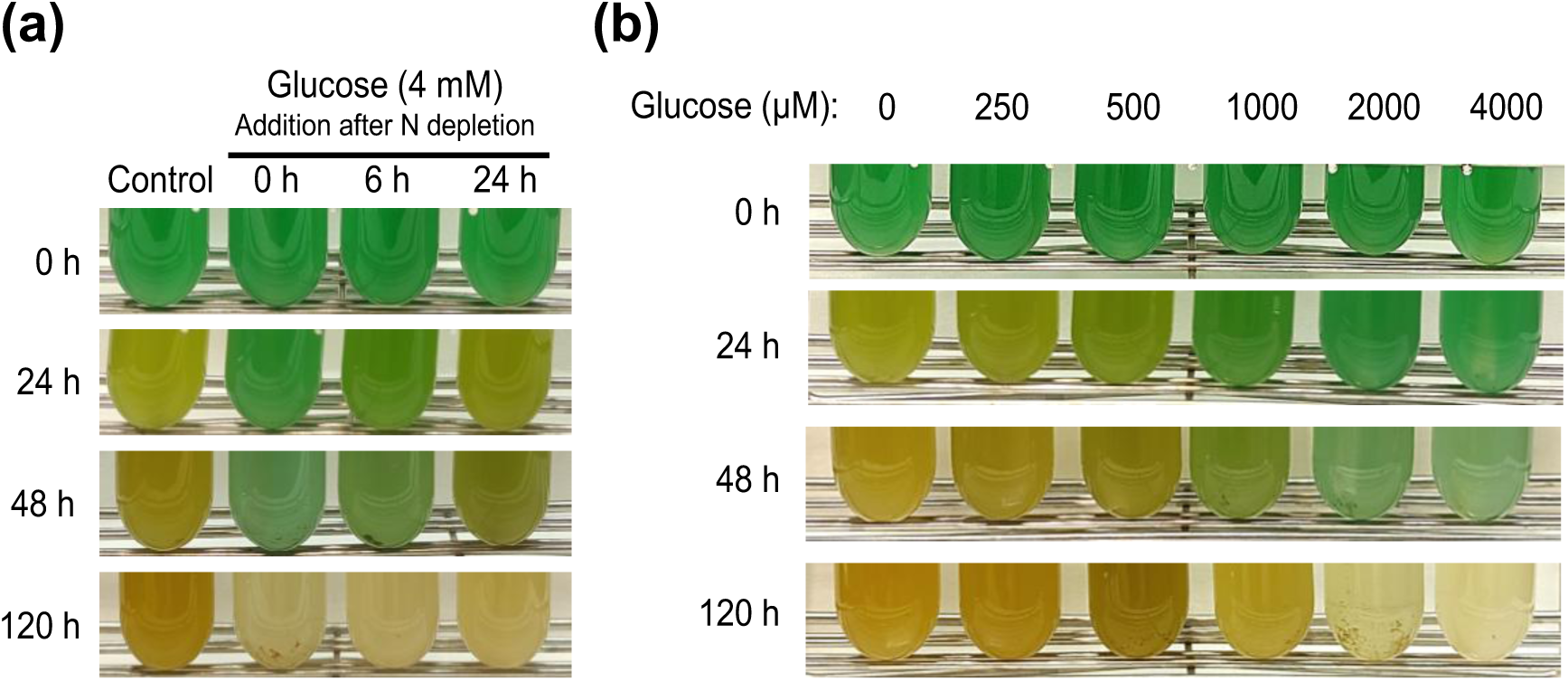
Effect of glucose concentration and time of addition on the bleaching process in response to nitrogen deprivation in WT *Synechocystis*. (a) Photographs of the growth of the WT strain cultured in nitrogen deplete medium (BG11_0_C) with addition of glucose at certain times after nitrogen deprivation. (b) Photographs of the growth of the WT strain cultured in BG11_0_C with different amounts of glucose added at the onset of nitrogen deprivation ranging from 250 to 4000 µM.

## Discussion

In photosynthetic organisms, maintaining balanced carbon pools is crucial to adapt to environmental changes while preventing electron transport dysregulation. *Synechocystis* can perform chlorosis (bleaching) under nitrogen deprivation or can growth photomixotrophically with glucose, both conditions requiring substantial metabolic adaptation and restructuring the photosynthetic machinery (Ortega-Martínez et al. 2023; Ortega-Martínez et al. 2024). Glycogen plays a critical role in these processes, acting as a metabolic buffer that coordinates source-sink relationships within the cell. In fact, nitrogen deficiency adaptation is characterized by an increase in the C/N ratio leading to accumulation of glycogen, partially supported by carbon recycled from degraded phycobiliproteins (Fig. 1c and (Hasunuma et al. 2013; Ortega-Martínez et al. 2023)). However, glucose supplementation under nitrogen deprivation further enhances the C/N ratio and doubles glycogen reserves within 6 hours (Fig. 1c), preventing PBS degradation and bleaching (Fig. 1a), in agreement with previous report (Elmorjani and Herdman 1987).

This suggests the combination of external glucose and PBS-derived carbon may exceed the buffering capacity of glycogen, thereby disrupting the chlorosis program. Consistent with this, we observed substantial alterations in carbon partitioning and accumulation of key metabolic intermediates (e.g., S7P, E4P, F6P, pyruvate, 2-OG, 6PG, malate, fumarate, succinate; Fig. 1e), indicating metabolic overflow. Such imbalances likely interfere with allosteric regulations and metabolic checkpoints essential for the progression of chlorosis as a nitrogen depletion response. This conclusion is supported by the non-bleaching phenotypes of mutants in enzymes such as alanine dehydrogenase or AGP, which also generate metabolic alterations (Lahmi et al. 2006; Carrieri et al. 2012; Gründel et al. 2012; Hickman et al. 2013; Ortega-Martínez et al. 2023).

Despite the lack of bleaching and the presence of a metabolic imbalance, the transcriptomic response to nitrogen deprivation in the presence of glucose was remarkably similar to that observed under nitrogen depleted conditions (Fig. 3). This response was consistent with earlier studies in *Synechocystis* during chlorosis (Krasikov et al. 2012; Matthias et al. 2014; Carrieri et al. 2017; Giner-Lamia et al. 2017; Esteves-Ferreira et al. 2018), where nitrogen deprivation induced genes involved in nitrogen uptake and transport systems, as well as *nblA* expression, while repressing transcripts for phycobiliproteins and carbon fixation pathways. This suggests that post-transcriptional or metabolic level regulation, rather than transcriptional regulation alone, contribute to the suppression of bleaching by glucose, independently of *nblA* expression. This conclusion is supported by non-bleaching mutants, such as the AGP (*glgC*) glycogen synthesis mutant (Hickman et al. 2013; Carrieri et al. 2017) or the mutant lacking the GntR-family regulator *sll1961* (Ozaki et al. 2007; Sato et al. 2008), expressing *nblA* during nitrogen deprivation but not bleaching. The observation that glucose addition halts bleaching even after the process has been initiated (Fig. 6) supports the existence of a superimposed mechanism of chlorosis control independent of the sensing and transcriptional response to nitrogen deficiency. Strikingly, the combined stress of nitrogen depletion and glucose supplementation in wild-type cells mirrors the behaviour of glycogen-deficient mutants (ΔAGP, ΔPGM) under these stresses separately, showing impaired viability, metabolic overflow and decreased photosynthetic activity (Ortega-Martínez et al. 2023; Ortega-Martínez et al. 2024). This suggests that chlorosis program might be directly influenced and disturbed by alterations in these processes, alongside the established role of NtcA transcription factor, nitrogen sensing and C/N ratio. Supporting this, our photosynthetic measurements of the nitrogen depleted WT culture supplemented with glucose showed a rapid decline in performance upon glucose addition during nitrogen deprivation (Fig. 4), similar to observations in glycogen mutants under nitrogen deficiency or glucose supplementation (Carrieri et al. 2012; Ortega-Martínez et al. 2023; Ortega-Martínez et al. 2024). Both oxygen evolution and Y(II) were already affected after three hours of glucose treatment, indicating that most PSII reaction centers are closed (Fig. 4a-d) and Q_A_ reduced (Fig. 4e and S6), with an expected reduced PQ pool. This photosynthetic condition agrees with the interpretation that nitrogen deficit acclimation requires a fully functional photosynthetic apparatus driven by actinic light, since PSII inhibition by DCMU or dark cultivation impedes the bleaching process (Salomon et al. 2013; Yoshihara and Kobayashi 2022).

An over reduced PQ pool could disrupt electron flow, prior to Cyt *b*_₆_*f*, via the RISE (reduction-induced suppression of electron flow) mechanism (Shimakawa et al. 2018) due to lack of free oxidized PQ available to participate in the Q cycle at Cyt *b*_6_*f*. However, overall data from different sources indicates that PQ pool is not governing the bleaching process. Firstly, the similarities in the data of photomixotrophic conditions in glycogen synthesis ΔAGP and ΔPGM strains (Ortega-Martínez et al. 2024) suggest that it is unlikely. Secondly, disrupting the OPP shunt by deleting the G6PDH enzyme (ΔG6PDH strain), and thus limiting the NADPH synthesis to the FNR reaction, did not ameliorate the impairments in the bleaching process caused by the glucose supplementation (Fig. 2). Thirdly, we cannot confirm the redox state of the PQ pool since estimations through qL and qP calculations (Fig. 4e and S6) do not provide a direct measurement; rather, they infer it from fluorescence parameters related to Q_A_ which is a known limitation of fluorescence-based proxies (Khorobrykh et al. 2020). This aligns with DCMU-treated cells, where Q_A_ is reduced but the PQ pool remains mostly oxidized due to blocked Q_B_ site within PSI, while fluorescence parameters qP and qL would indicate a reduction of PQ pool.

Further insight came from PSI redox measurements. Glucose-supplemented, nitrogen-depleted cultures showed a decline in Y(I), increased donor-side limitation [Y(ND)] and P700⁺ accumulation during actinic illumination (Fig. 5). This aligns with decreased electron flow from PSII to PSI and suggests that a sufficient electron transfer from PSI may be a prerequisite to proceed chlorosis, while an oxidized P700/P700^+^ could act as a redox signal preventing bleaching. Nevertheless, under nitrogen deplete conditions, the quantitatively strong photosynthetic electron sink from Fd, that suppose nitrate reduction to ammonia, is not active. This scenario contributes to reduction of the Fd pool and thus, diminished the oxidation of its electron donor PSI (P700), becoming a key component to trigger the response to nitrogen shortage. Similar conclusions on the role of a reduced P700 in bleaching process were reached on *Synechochoccus*, where promoting the oxidation of the PETC by adding nitrate (and thus consuming reduced Fd) prevented the induction of chlorosis and *nblA* expression caused by sub-toxic MSX treatment (glutamine synthetase inhibitor) in the presence of ammonia (Klotz et al. 2015).

This assumption that bleaching signal is governed by a reduced P700 threshold, and not by other PETC components like the redox state of PQ pool, is emphasized by DCMU or DBMIB treatments: Both inhibitors of photosynthesis prevent the beaching process (Yoshihara and Kobayashi 2022) disrupting electron transfer at the level of the PQ pool, causing its overall oxidized state (DCMU) or complete reduction (DBMIB).

Together, our results demonstrate that under nitrogen deprivation and glucose supplementation, the photosynthetic electron transport chain is disrupted at the PSII acceptor side (between Q_A_ and Q_B_). This leads to lack of electron flow to and from PSI. This electron flow bottleneck underlies the failure to proceed with bleaching and highlights a redox-controlled checkpoint in the nitrogen starvation response of *Synechocystis*. Furthermore, these results emphasize the thigh metabolic-photosynthetic control of photosynthetic organisms.

## Author contributions

P.O.M., F.J.F. and S.D.T conceived the study together. P.O.M carried out the experimental work, and was responsible for data analysis, visualization, and drafting the original manuscript. J.G.L. contributed to formal analysis and data visualization. L.T.W. provided essential resources and support for data analysis and visualization. M.I.M. and F.J.F. oversaw project administration and secured funding. S.D.T. supervised the project and contributed to data interpretation and visualization. All authors contributed to reviewing and editing the final manuscript.

## Acknowledgements

This work was funded by grants PID2019-104513GB-I00 and PID2022-138317NB-I00 both financed by MCIU/AEI/10.13039/501100011033/ ‘FEDER Una manera de hacer Europa’ and by Junta de Andalucía, Group BIO-0284 to FJF.

## Data availability statement

The RNA sequencing data used for Fig. 3 and Tables S2-4 are available in NCBI under accession number PRJNA1292945. Other data supporting the findings of this study are available in the manuscript or supplementary materials

