## Supplemental Figures for "A photosynthetic checkpoint halts *Synechocystis* sp. PCC 6803 response to nitrogen deprivation during glucose-mediated metabolic imbalance"

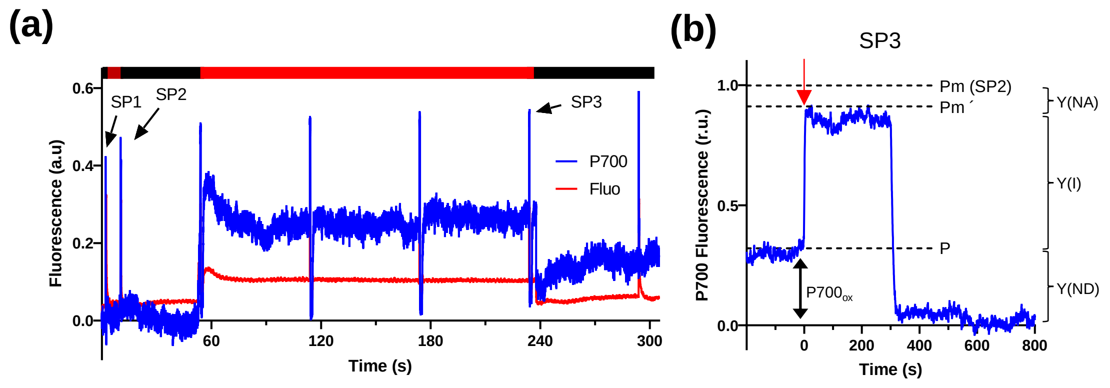


**Figure S1. Example of a P700 redox kinetic**.

(a) A representative induction curve (IC) in a Dual-PAM-100 for the simultaneous measurements of Chl fluorescence and redox kinetics of P700. The induction curve starts with a saturating pulse SP in darkness (SP1), where P700 is reduced. That is followed by the determination of maximum photooxidisable (Pm), achieved by a SP under far-red illumination (SP2), which preferentially excites PSI. This value is used in combination with the steady-state P700 oxidation level under actinic light (P) and maximum P700 oxidation level (Pm´) under a SP during actinic light (SP3) to calculate photosystem I quantum yields. (See figure 5). (b) Detail of a SP applied during actinic light (SP3) and description of parameters calculated, the effective quantum yield of PSI [Y(I)], and the non-photochemical energy dissipation due to donor-side [Y(ND)] and acceptor-side [Y(NA)] limitations. The red arrow denotes the onset of the 300 ms saturation pulse at 5000 µmol photons $m^{-2}$ $s^{-1}$.

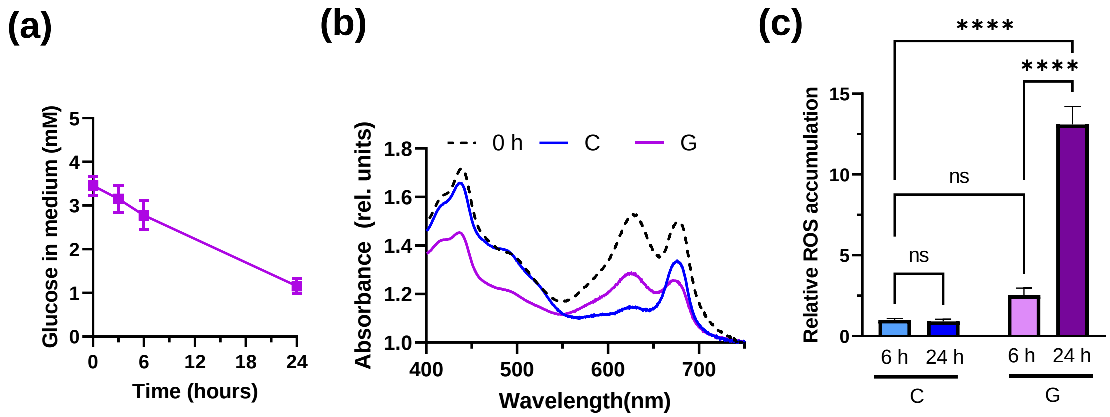


**Figure S2. Consumption of glucose during nitrogen depletion and its effect on whole cell absorption profile, and reactive oxygen species production**

WT cells grown in nitrogen-replete medium were harvested and resuspended in nitrogen-free ($BG11_0C$) medium at 1 $OD_{750}$. The culture was divided and grown under control (C) conditions or supplemented with 4 mM of glucose (G). (a) Glucose concentration profile in the media. (b) Representative whole cell spectra before (0 h) and 24 hours after culture under control conditions or supplemented with 4 mM of glucose. (c) Relative quantification of cellular reactive oxygen species (ROS) measured as 2',7'-Dichlorofluorescein fluorescence in cells cultured 6 and 24 hours in C and G conditions compared with levels of cells growing in nitrogen replete conditions. Data in (a) and (c) represent the mean±SEM of 4 independent biological replicates. Statistical significance was denoted as ****, P<0.0001; and ns, no significance (two-way ANOVA).

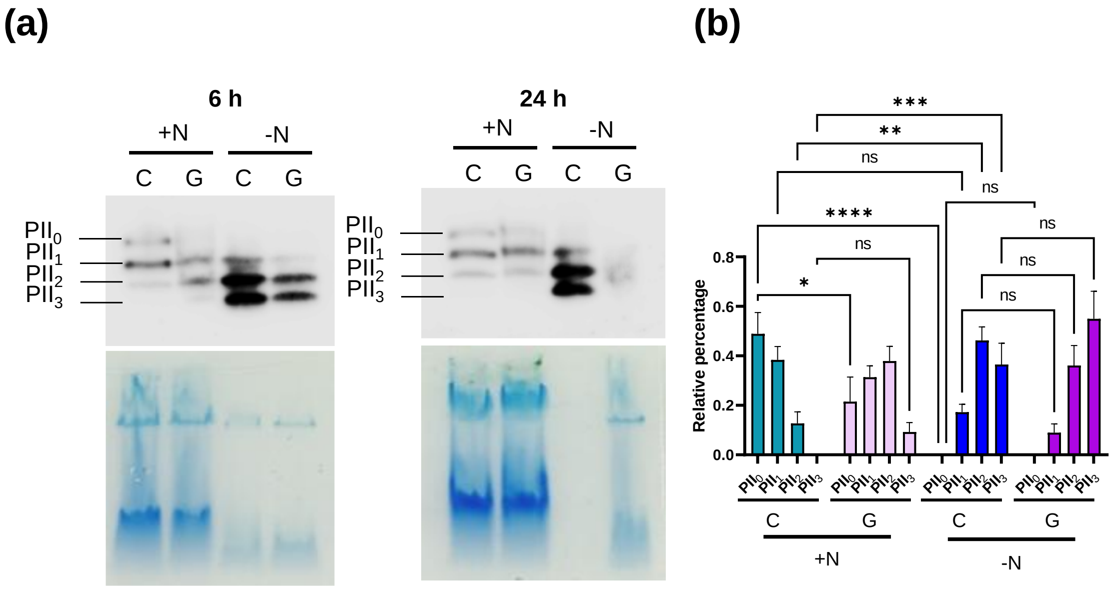


**Figure S3. Phosphorilation state of PII in response to nitrogen deprivation and glucose supplementation**

Native-PAGE analysis of soluble extracts of control (C) and glucose supplemented (G) cells after 6 and 24 hours in the presence (+N) or absence (+N) of nitrate. (a) Representative image of the immunochemoluminiscence against P-II protein (polyclonal antibody; 1:4000) and of the unstained polyacrylamide gels after electrophoresis of a total of 4 independent biological replicates is shown. (b) Quantification of phosphorylated PII bands after 6 hours of treatment. Data represent the mean±SEM of 4 independent biological replicates. Statistical significance was denoted as *, P<0.05; **, P<0.01; ***, P<0.001; ****, P<0.0001; ns, no significance (two-way ANOVA).

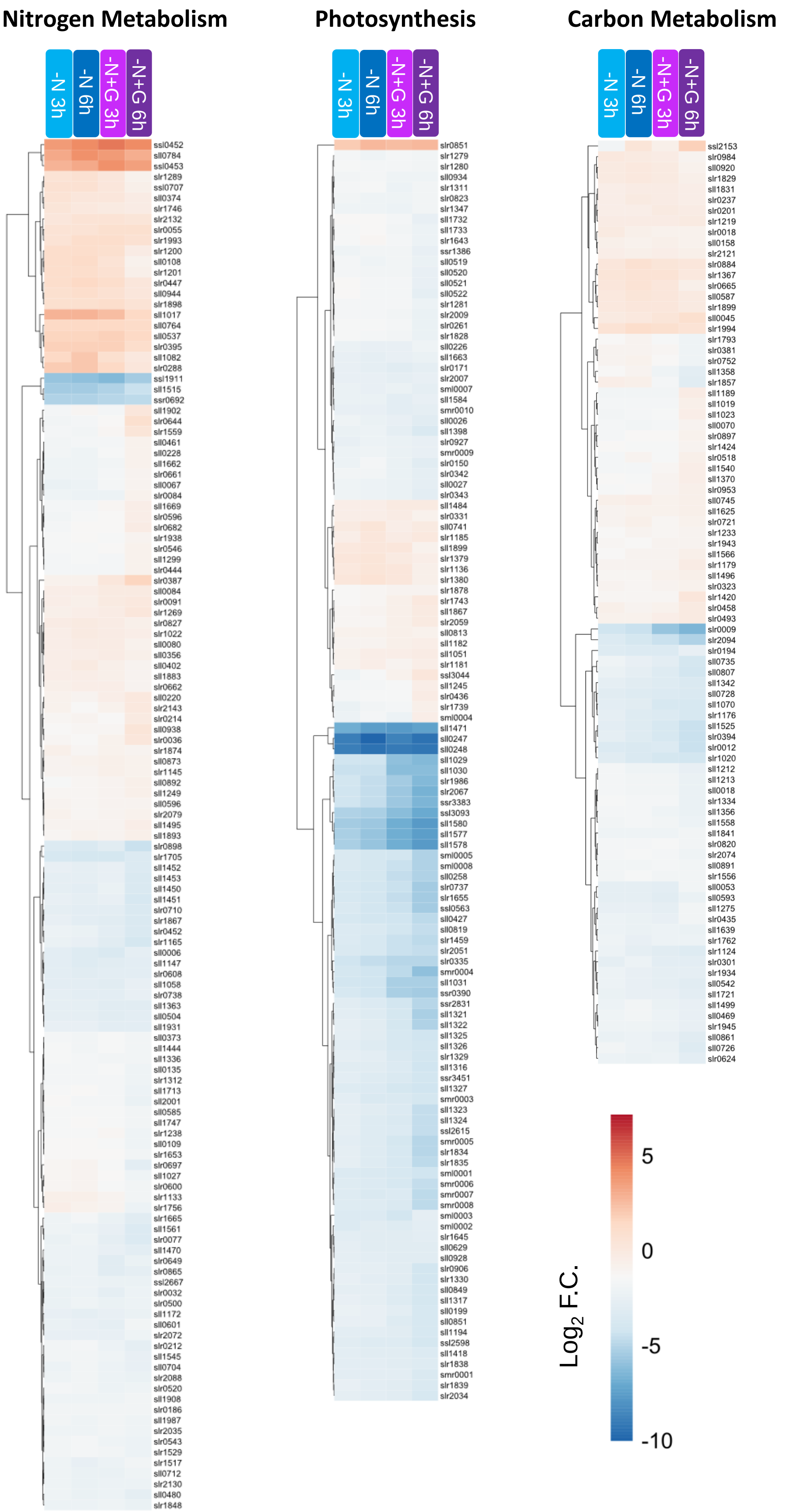
Nitrogen Metabolism
Photosynthesis
Carbon Metabolism
-N 3h
-N 6h
-N+G 3h
-N+G 6h
ssl0452
sll0784
ssl0453
slr1289
ssl0707
sll0374
slr1746
slr2132
slr0055
slr1993
slr1200
sll0108
slr1201
slr0447
sll0944
slr1898
sll1017
sll0764
sll0537
slr0395
sll1082
slr0288
ssl1911
sll1515
ssr0692
sll1902
slr0644
slr1559
sll0461
sll0228
sll1662
slr0661
sll0067
slr0084
sll1669
slr0596
slr0682
slr1938
slr0546
sll1299
slr0444
slr0387
sll0084
slr0091
slr1269
slr0827
slr1022
sll0080
sll0356
slr0402
sll1883
slr0662
sll0220
slr2143
slr0214
sll0938
slr0036
slr1874
sll0873
slr1145
sll0892
sll1249
sll0596
slr2079
sll1495
sll1893
slr0898
slr1705
sll1452
sll1453
sll1450
sll1451
slr0710
slr1867
slr0452
slr1165
sll0006
sll1147
slr0608
sll1058
slr0738
sll1363
sll0504
slr1931
sll0373
sll1444
sll1336
sll0135
slr1312
sll1713
sll2001
sll0585
sll1747
slr1238
sll0109
slr1653
slr0697
sll1027
slr0600
slr1133
slr1756
slr1665
sll1561
slr0077
sll1470
slr0649
slr0865
ssl2667
slr0032
slr0500
sll1172
sll0601
slr2072
slr0212
sll1545
sll0704
slr2088
slr0520
sll1908
slr0186
sll1987
slr2035
slr0543
slr1529
slr1517
sll0712
slr2130
sll0480
slr1848
slr0851
slr1279
slr1280
sll0934
slr1311
slr0823
slr1347
sll1732
sll1733
slr1643
ssr1386
sll0519
sll0520
sll0521
sll0522
slr1281
slr2009
slr0261
slr1828
sll0226
sll1663
slr0171
slr2007
sml0007
sll1584
smr0010
sll0026
sll1398
slr0927
smr0009
slr0150
slr0342
sll0027
slr0343
sll1484
slr0331
sll0741
slr1185
sll1899
slr1379
slr1136
slr1380
slr1878
slr1743
sll1867
slr2059
sll0813
sll1182
sll1051
slr1181
ssl3044
sll1245
slr0436
slr1739
sml0004
sll1471
sll0247
sll0248
sll1029
sll1030
slr1986
slr2067
ssr3383
ssl3093
sll1580
sll1577
sll1578
sml0005
sml0008
sll0258
slr0737
slr1655
ssl0563
sll0427
sll0819
slr1459
slr2051
slr0335
smr0004
sll1031
ssr0390
ssr2831
sll1321
sll1322
slr1325
sll1326
slr1329
sll1316
ssr3451
sll1327
smr0003
sll1323
sll1324
ssl2615
smr0005
slr1834
slr1835
sml0001
smr0006
smr0007
smr0008
sml0003
sml0002
slr1645
sll0629
sll0928
slr0906
slr1330
sll0849
sll1317
sll0199
sll0851
sll1194
ssl2598
sll1418
slr1838
smr0001
slr1839
slr2034
ssl2153
slr0984
sll0920
slr1829
sll1831
slr0237
slr0201
slr1219
slr0018
sll0158
slr2121
slr0884
slr1367
slr0665
sll0587
slr1899
sll0045
slr1994
slr1793
slr0381
slr0752
sll1358
slr1857
sll1189
sll1019
sll1023
sll0070
slr0897
slr1424
slr0518
sll1540
sll1370
slr0953
sll0745
sll1625
slr0721
slr1233
slr1943
sll1566
slr1179
sll1496
slr0323
slr1420
slr0458
slr0493
slr0009
slr2094
slr0194
sll0735
sll0807
sll1342
sll0728
sll1070
slr1176
sll1525
slr0394
slr0012
slr1020
sll1212
sll1213
sll0018
slr1334
sll1356
sll1558
sll1841
slr0820
slr2074
sll0891
slr1556
sll0053
sll0593
sll1275
slr0435
sll1639
slr1762
slr1124
slr0301
slr1934
sll0542
sll1721
sll1499
sll0469
slr1945
sll0861
sll0726
slr0624
5
0
-5
-10
$Log_2$ F.C.

**Figure S4. Effect of glucose supplementation on the transcriptomic response of *Synechocystis* to nitrogen deprivation in genes related to carbon metabolism, photosynthesis, respiration, and nitrogen assimilation.**

Heatmap showing the $log_2$ fold-change of significantly differentially expressed genes (adj. *p-value* < 0.05) related to carbon metabolism, photosynthesis, respiration, and nitrogen assimilation after 3 and 6 hours under nitrogen deprivation (–N) and nitrogen deprivation with glucose supplementation (–N+G), compared to nitrogen-replete controls (0h). Comparisons shown: *–N 3h vs 0h, –N 6h vs 0h, –N+G 3h vs 0h, and –N+G 6h vs 0h*. $Log_2$ fold-change values are represented using a colour scale: red indicates upregulation and blue indicates downregulation.

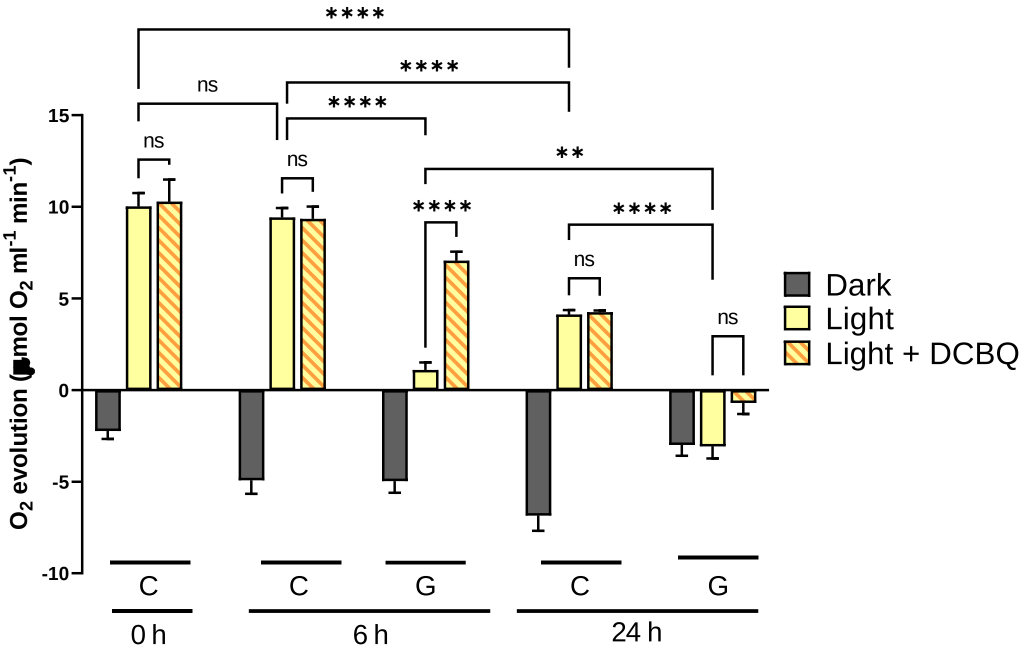


**Figure S5. Effect of DCBQ on oxygen evolution during nitrogen depletion and glucose supplementation.**

WT cells grown in nitrogen replete medium (BG11C) were harvested and resuspended in nitrogen-free medium ($BG11_0C$) at 1 $OD_{750}$. The culture was divided and grown under control (C) conditions or supplemented with 4 mM of glucose (G). The oxygen evolution rates of 2 mL of cultures at 6 and 24 hours after nitrogen deprivation were recorded on a Clark-type electrode with an illumination of 50 µmol photons $m^{-2}$ $s^{-1}$ and adding to the chamber a fresh mix of DCBQ (0,25 mM) and ferricyanide (2,5 mM). Data represent the mean±SEM of 4 independent biological replicates. Statistical significance was denoted as **, P<0.01; ****, P<0.0001; and ns, no significance (two-way ANOVA).

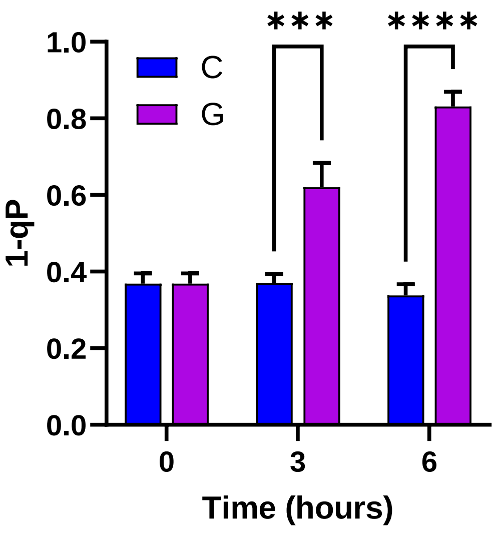


**Figure S6. Chlorophyll fluorescence estimation of 1-qP during nitrogen depletion and glucose supplementation.**

Estimations of 1-qP obtained from the traces in Figure 4. Data represent the mean±SEM of 5 independent biological replicates. Statistical significance was denoted as ***, P< 0.001 and ****, P<0.0001 (two-way ANOVA).

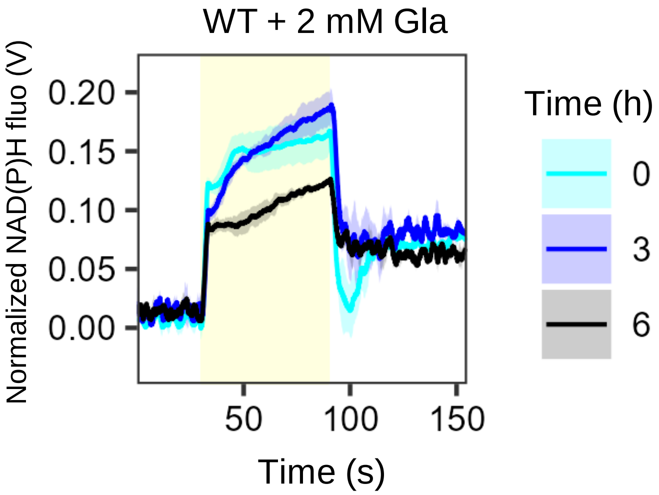


**Figure S7. Effect of Gla treatment in NAD(P)H fluorescence light-dependent kinetics in *Synechocystis***

WT cells grown in nitrogen replete medium (BG11C) were harvested and resuspended in nitrogen-free medium ($BG11_0C$) at 1 $OD_{750}$ with 2 mM of glycolaldehyde (Gla). NAD(P)H fluorescence light-dependent kinetics was registered at 0, 3 and 6 hours after nitrogen depletion and . Data represent the mean±SEM of 4 independent biological replicates.
