## Supplemental Material and Methods for "A photosynthetic checkpoint halts *Synechocystis* sp. PCC 6803 response to nitrogen deprivation during glucose-mediated metabolic imbalance"

### Protein extraction and immunoblotting

Fractions of 15 ml of cultures were harvested at six and 24 hours by centrifugation (5000 g 10 min 4º C) and storage frozen at -20º C. Frozen pellets were resuspended in 200 µl 50 mM Tris-HCl pH 7.5, 50 mM NaCl and 1 mM phenylmethylsulfonyl fluoride (PMFS) with glass beads (equivalent volume to 150-200 µL). Cell lysates were obtained after mechanical disruption by automatic disruption (at 6m/s for 30s in a FastPrep-24 5G; MP Biomedicals). Samples were centrifugated at 4 ºC for 20 min at 15000 g and 150 µl of supernatant, containing soluble proteins, was collected in a new tube and mixed with 50 µl of loading buffer without reducing agents or SDS. In general, 20 µl of each sample was resolved by Native-PAGE (15% acrylamide/bis-acrylamide). For –N at six hours samples and for C –N at 24 hours, 5- and 10-fold dilutions were loaded. After electrophoresis, proteins were transferred to a polyvinylidene difluoride membrane (Immobilon-P; Millipore). Blocking, probing and detection was performed as in (Ortega-Martínez et al. 2023) using primary antibody against P-II protein (1:4000) diluted in blocking solution.
